# Absence of cardiolipin from the outer leaflet of a mitochondrial inner membrane mimic restricts Opa1-mediated fusion

**DOI:** 10.1101/2021.09.01.458556

**Authors:** Yifan Ge, Sivakumar Boopathy, Tran H Nguyen, Camila Makhlouta Lugo, Luke H. Chao

**Affiliations:** Department of Molecular Biology, Massachusetts General Hospital, Boston, Massachusetts, U.S.A, 02114; Department of Genetics, Harvard Medical School, Boston, Massachusetts, U.S.A.

## Abstract

Cardiolipin is a tetra-acylated di-phosphatidylglycerol lipid enriched in the matrix-facing (inner) leaflet of the mitochondrial inner membrane. Cardiolipin plays an important role in regulating mitochondria function and dynamics. Yet, the mechanisms connecting cardiolipin distribution and mitochondrial protein function remain indirect. In our previous work, we established an *in vitro* system reconstituting mitochondrial inner membrane fusion mediated by Opa1. We found that the long form of Opa1 (l-Opa1) works together with the proteolytically processed short form (s-Opa1) to mediate fast and efficient membrane fusion. Here, we extend our reconstitution system to generate supported lipid bilayers with asymmetric cardiolipin distribution. Using this system, we find the presence of cardiolipin on the inter-membrane space-facing (outer) leaflet is important for membrane tethering and fusion. We discuss how the presence of cardiolipin in this leaflet may influence protein and membrane properties, and future applications for this approach.

## Introduction

Mitochondria are important eukaryotic organelles that regulate cell metabolism, signaling and death (McBride et al., 2006;Giacomello et al., 2020). Mitochondria feature a unique double membrane structure. The outer mitochondrial membrane (OMM) hosts machinery for lipid and protein transport, while the inner mitochondrial membrane (IMM), with its cristae invaginations, is the site for ATP synthesis and respiratory function (Kroemer et al., 2007)(Cogliati et al., 2016;Kondadi et al., 2020). Mitochondrial membranes are dynamic, undergoing fusion and fission to distribute metabolites and buffer mutation in mitochondrial DNA, while changing morphology in response to physiological conditions (Chan, 2020;Gao and Hu, 2021).

While phospholipid composition is recognized as a key regulator of membrane properties, the functional roles of phospholipid distribution are challenging to investigate (Harayama and Riezman, 2018). The mitochondrial membrane is distinct in composition from other endomembrane systems, and comprises cardiolipin (CL), phosphocholine (PC), and phosphatidylethanolamine (PE) (Horvath and Daum, 2013). Other lipid species that may play important roles include phosphatidylinositol (PI), phosphatidylserine (PS), phosphatidylglycerol (PG), phosphatidic acid (PA), lysophospholipids, sterols, sphingomyelin and low levels of cholesterol (CH) (Horvath and Daum, 2013).

In eukaryotes, CL is uniquely found in the mitochondria. Consistent with an endosymbiotic origin, CL is found in bacteria (Friedman and Nunnari, 2014). CL comprises ∼20% of total mitochondrial phospholipid content. In the OMM, PC and PE are major membrane components. In contrast, PE and CL makes up nearly 50% of the total phospholipid in the IMM (Horvath and Daum, 2013; Oemer et al., 2018; Oemer et al., 2020). Electron paramagnetic resonance spectroscopy studies of yeast mitochondria show CL enrichment in the inner leaflet of the IMM (Petit et al., 1994). An inner-leaflet IMM distribution of CL was also shown using fluorescent probes that specifically target CL (Krebs et al., 1979; Harb et al., 1981; Petit et al., 1994). These observations are consistent with data indicating CL is synthesized from PA in the matrix-proximal face of the IMM (Schlame, 2008). While the distribution between leaflets may be dynamic, data suggest CL also comprises ∼5-15% of the total lipids in the inter-membrane space (IMS)-facing (outer) leaflet (Harb et al., 1981;Gallet et al., 1997).

CL has four acyl chains, forming a unique conical structure expected to have high flexibility. This structure classifies CL as a ‘non-structural’ phospholipid that tends to introduce a hexagonal phase in mixtures (Kagan et al., 2014). The high concentration of PE and CL in the IMM is expected to alter membrane tension (Traïkia et al., 2002), which influences membrane bending. Differential stress between leaflets can change membrane elasticity, stiffness and lateral pressure (Hossein and Deserno, 2020). These changes can favor incorporation of transmembrane proteins that induce curvature, association of peripheral membrane proteins, or self-assembly of proteins on either leaflet (Wilson et al., 2019;Lorent et al., 2020).

In addition to inter-leaflet asymmetry, lipids can exhibit planar segregation (Doktorova et al., 2020; Lorent et al., 2020). Lateral heterogeneity in the plasma membrane has been more extensively studied (Lorent et al., 2020). For example, model plasma membrane experiments show planar segregation coupled to inter-leaflet asymmetry can affect lipid packing and hydrophobic thickness, to sequester transmembrane proteins (Hussain et al., 2013). Similar physical consequences of planar segregation may be at play in the IMM. Although lateral segration and principles of IMM organization are less well studied in well-controlled biophysical systems, cellular experiments suggest membrane heterogeneity may play a role in organizing respiratory chain super-complexes (Dudek, 2017).

Theory and computation have been applied to investigate the physical consequences of heterogeneity in CL distribution (Lemmin et al., 2013;Boyd et al., 2017;Elias-Wolff et al., 2019;Wilson et al., 2019). Not surprisingly, CL distribution is predicted to strongly shape the electrostatic potential of membrane surface, with cation interactions observed to introduce lateral CL heterogeneity in simulations. (Lemmin et al., 2013). Boyd and coworkers demonstrated concentration-dependent bending and buckling of CL bilayers, with CL mediating regions of high curvature, and CL concentrating in the negatively curved regions (Boyd et al., 2017). This finding was supported by work from Elias-Wolf and coworkers, which also found CL inter-leaflet asymmetry contributes to membrane buckling (Elias-Wolff et al., 2019).

CL distribution can change in response to environmental stimuli, and has important consequences in disease (Gallet et al., 1997). Recent studies show CL can be externalized by pro-mitophagy stimuli in primary cortical neurons and in cellular models of Parkinson’s Disease (Chu et al., 2013). Externalization of CL to the outer leaflet of the OMM activates binding to microtubule-associated-protein-1 light chain 3, which localizes damaged mitochondria to autophagosomes (Chu et al., 2013; Ryan et al., 2018). CL externalization is recognized by macrophages to cause inflammation (Pizzuto and Pelegrin, 2020). These processes are associated with altered fission and fusion dynamics and altered mitochondrial morphology (Kordower et al., 2008; Van Laar and Berman, 2009). Many mitochondrial proteins depend on CL for activity (Ban et al., 2010; Zhang et al., 2020), yet the role of asymmetric lipid distribution on protein activity has been technically challenging to investigate.

In our previous work, we fabricated a model membrane system that mimicked the IMM in composition. We reconstituted Opa1 (the IMM fusogen) into a supported bilayer and noticed CL played a key role in several stages of membrane fusion (in particular, GTP-dependent membrane tethering). In our previous study, we focused on understanding how the full-length ‘long form’ of Opa1 (l-Opa1) worked together with a proteolytically processed ‘short form’ (s-Opa1) to regulate full fusion. We showed that while l-Opa1 can induce low levels of membrane fusion, both s-Opa1 and l-Opa1 are required for fast and efficient fusion. These experiments were performed under conditions containing symmetric lipid composition in the bilayer, with both leaflets containing 20 % CL. In these previous experiments, we found that when the model membranes lacked CL entirely, fusion was rare.

Here, we present new data investigating Opa1 activity under asymmetric bilayer conditions. We take advantage of our synthetic model-membrane system to generate supported lipid bilayers with asymmetric CL distribution. Our new results show CL in the outer leaflet (facing the IMS) plays an important role in Opa1-mediated bilayer tethering and pore opening. These data implicate CL-protein interactions during specific steps of membrane fusion. We discuss how changes in bilayer composition may influence membrane properties and protein-protein interactions.

## Materials and Methods

### Preparation of polymer-tethered asymmetric bilayers and reconstitution of l-Opa1 into asymmetric bilayers

Polymer-tethered lipid bilayers were fabricated using mixtures of synthesized lipid reagents, including 1,2-dioleoyl-sn-glycero-2-phosphocholine, (DOPC); 1,2-dioleoyl-sn-glycero-2-phosphoethannolamine-N-[metoxy(polyethylene glycerol)-2000] (DOPE-PEG2000), L-α-phoshphatidylinositol (liver PI) and 1’,3’-bis[1,2-dioleoyl-sn-glycero-3-phospho]-glycerol (18:1 cardiolipin) from Avanti Polar Lipids (AL, USA). Inter-leaflet asymmetric bilayers were assembled in a layer-by-layer manner, using a combination of Langmuir-Blodgett dipping and Langmuir-Schaefer transfer. Bilayers were fabricated on a glass substrate (25 mm diameter cover glass, Fisher Scientific) as previously described (Ge et al., 2020b). For all fusion experiments, the bottom leaflet included 7 %(mol) liver PI, 20 %(mol) CL, 20 %(mol) DOPE, 0.2 %(mol) Cy5-PE and 5 %(mol) DOPE-PEG2000. For fluorescence anisotropy analysis, the bottom leaflet contained 0.2 %(mol) Top-fluor CL (Avanti Polar Lipids), which served as a reporter for CL distribution and to evaluate lipid flip-flop. The lipid mixture was spread carefully on top of the air-water interface on a Langmuir-Blodgett trough (KSV-NIMA, NY, USA). A film was applied at a pressure of 37 mN/m and kept for 30 min., then transferred to the glass substrate, forming the bottom leaflet of the bilayer. The top leaflet was fabricated using a Langmuir-Schaefer transfer method following fabrication of the LB layer, as previously described (Ge et al., 2020b). For l-Opa1 reconstitution experiments, the top leaflet (mimicking the outer, IMS-facing side) of the asymmetric bilayer was completely CL free, whereas the bottom leaflet (facing the inner, matrix side) contains 20% CL. TopFluor CL was used to evaluate bilayer anisotropy. Texas Red was used to monitor hemifusion kinetics. For all bilayers, the lipid mixture of the bottom leaflet was spread on top of the air-water interface, while pressure was kept at 37 mN/m for 30 mins before dipping. The bottom leaflet was prepared at 30 mN/m pressured for 30 mins before Schaefer transfer and assembly into an asymmetric bilayer. This helped maintain a similar area/molecule ratio in both top and bottom leaflets (Ge et al., 2020b).

### Purification and reconstitution of Opa1 in asymmetric bilayers

l- and s-Opa1 were purified following a protocol reported previously (Ge et al., 2020a;Ge et al., 2020b). Human l-Opa1 (isoform 1) and s-Opa1 with Twin-strep-tag at N-terminus and deca-histindine tag at C-terminus (GenScript, NJ, USA) was expressed in *Pichia pastoris* strain SMD1163. The cells were harvested and milled using SPEX 6875D Freezer/Mill. The milled powder was resuspended in buffer containing 50 mM sodium phosphate, 300 mM NaCl, 1 mM 2-mercaptoethanol, pH 7.5, with benzonase nuclease and protease inhibitors. The membrane fraction was collected by ultra-centrifugation at 235,000 xg for 45 min. at 4 °C, then incubated for 1 hour in the same resuspension buffer supplemented with 2 % DDM (Anatrace, OH, USA) and 0.1 mg/ml cardiolipin. Membrane fraction solubilized in DDM containing buffer was ultracentrifugated at 100,000 x g for 1 hour at 4 °C. The supernatant extract containing l-Opa1 was loaded onto a Ni-NTA column (Biorad, CA, USA), washed with 50 mM sodium phosphate, 350 mM NaCl, 1 mM 2-mercaptoethanol, 1 mM DDM, 0.025 mg/ml 18:1 CL, pH 7.5, with 25 mM imidazole followed with 100 mM imidazole. The bound Opa1 was eluted with the above buffer containing 500 mM imidazole. The elution was buffer exchanged into 100 mM Tris-HCl, 150 mM NaCl, 1 mM EDTA, 1 mM 2-mercaptoethanol, 0.15 mM DDM, 0.025 mg/ml 18:1 CL, pH 8.0. The C-terminal His-tag was then cleaved by treatment with TEV protease and applied to Ni-NTA and Strep-Tactin XT Superflow column (IBA life Sciences, Göttingen, Germany) attached in tandem and washed. The Strep-Tactin XT column was detached and the protein eluted with buffer containing 50 mM biotin. The elution fraction was concentrated and subjected to size exclusion chromography in buffer with 25 mM Bis-Tris Propane, 100 mM NaCl, 1 mM TCEP, 0.025 mg/ml 18:1 CL, pH 7.5.

l-Opa1 was reconstituted into bilayers by mixing purified l-Opa1 with a surfactant cocktail containing DDM and OG, applied as reconstitution reagents, at a concentration of 1.2 nM and 1.1 nM, respectively. l-Opa1 was reconstituted to total amount of 1.3 × 10^−12^ mol (protein:lipid 1:100000). The protein was incubated for 2 hrs in surfactant at room temperature. Extra surfactant was then removed using Bio-Beads SM2 (Bio-Rad, CA, USA), at a concentration of 10 μg/ml.

### Preparation of liposomes and proteoliposomes

Calcein encapsulated liposomes containing 7 %(mol), PI 20 %(mol), CL, 20 PE % (mol), 0.2 %(mol), TexasRed-PE, and DOPC 52.8 %(mol), were prepared as previously described (Ge et al., 2020a; Ge et al., 2020b). Lipids dissolved chloroform were mixed in the approproiate ratio and dried under nitrogen flow for 20-30 mins. The resulting lipid film was hydrated with calcein containing Bis-Tris buffer (50 mM calcein, 25 mM Bis-Tris, 150 mM NaCl, pH 7.4) at 70 °C for 30 mins following vigorous agitation. The resulting vesicles were homogenized using a mini-extruder using 200 nm polycarbonate membrane for 15-20 times extrusion to prepare large unilamellar vesicles (LUVs).

l-Opa1 was reconstituted in the LUVs at a molar ratio of 1:5000. Purified l-Opa1 was added to the LUV suspensions with DDM (final concentration 0.1 μM) and incubated for 2 hours at 4 °C. After reconstitution, the sample was dialyzed against calcein buffer using a 3.5 kDa dialysis cassette at 4 °C overnight. Calcein that was not encapsulated in the liposomes were removed using a PD10 desalting column.

### TIRF microscopy and fluorescence anisotropy analysis

Fluorescence images were taken using a Vector TIRF system (Intelligent Imaging Innovation, Inc, Denver, CO, USA) equipped with a W-view Gemini system (Hamamatsu photonics, Bridgewater, NJ). TIRF images were acquired using a 100X oil-immersion TIRF objective (Zeiss, N.A 1.4). Fluorescent images were taken using a Prime 95B scientific CMOS camera kept at -10 °C (Photometrics), recorded at a frame rate of 100 ms.

To validate bilayer asymmetry, fluorescence anisotropy was measured using a modified light path embedded in the microscopy. Under this experimental setting, the 488 nm excitation laser was polarized through a fiber switch system, equipped with internalized polarizer (Intelligent Imaging Innovation, Inc, Denver, CO, USA). The p- and s-emission were separated using a polarized beam splitter (532 nm, Thorlabs) inserted in the Gemini system. As a result, p-emission and s-emission was separated onto two halves of the camera chip. To ensure a reliable emission result, the p-polarized emission and s-polarized emission was adjusted to be similar intensity and was determined using a photometer. The fluorescence anisotropy of dye-labeled bilayers in both symmetric and asymmetric bilayers was determined by the following formula:

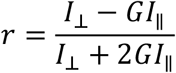

Where G represents the correction factor (added to compensate for uneven light distribution caused by the polarized emission beam splitter optics, and potential depolarization effects caused by the high NA objective). The G factor was determined using a sample solution containing 1 mM calcein dissolved in glycerol, where the fluorescence anisotropy was well defined. The G factor was evaluated for each pixel of the image before every measurement, and the final *r* of each pixel was obtained and averaged for the lipid bilayer samples to eliminate potential effects of inner leaflet dye aggregation.

For fusion and membrane tethering experiments, a 543 nm laser was used to analyze TexasRed-PE signal in liposomes and proteoliposomes and to observe lipid demixing. The bilayer was labeled with Cy5 (excited by a 633 nm laser), that reported on homogeneity of the bilayer. A 488 nm laser was applied for recording content release of the encapsulated calcein. Time-lapse images were taken at a frequency of 100 ms to record fusion dynamics.

## Results

### Bilayers with asymmetric cardiolipin distribution can be generated through layer-by-layer assembly

We generated a PEG polymer-tethered lipid bilayer with asymmetric CL distribution that mimics the leaflet distribution found in the mitochondrial inner membrane. Our approach was inspired by methods successfully used to study the leaflet asymmetry of cholesterol in the plasma membrane (Hussain et al., 2013). These methods have been used to study phase separation in planar lipid bilayers and liposomes (Hussain et al., 2013; Doktorova et al., 2018; London, 2019). Our control symmetric bilayers comprised 15 %(mol) CL on both top and bottom leaflets (with 0.1 % CL in both leaflets replaced by Top-Fluor CL) (Fig 1A). Control asymmetric bilayers comprised 10 %(mol) CL on the top leaflet and 20 %(mol) CL in the bottom leaflet (Top-Fluor CL only labeling the bottom leaflet) (Fig 1D). We noticed symmetric bilayers fabricated in our previous work had to be used immediately, as they acquired defects within 24 hours (Ge et al., 2020b). We did not observe this issue with our asymmetric bilayers, which were stable for at least 48 hours.

**Fig. 1.**
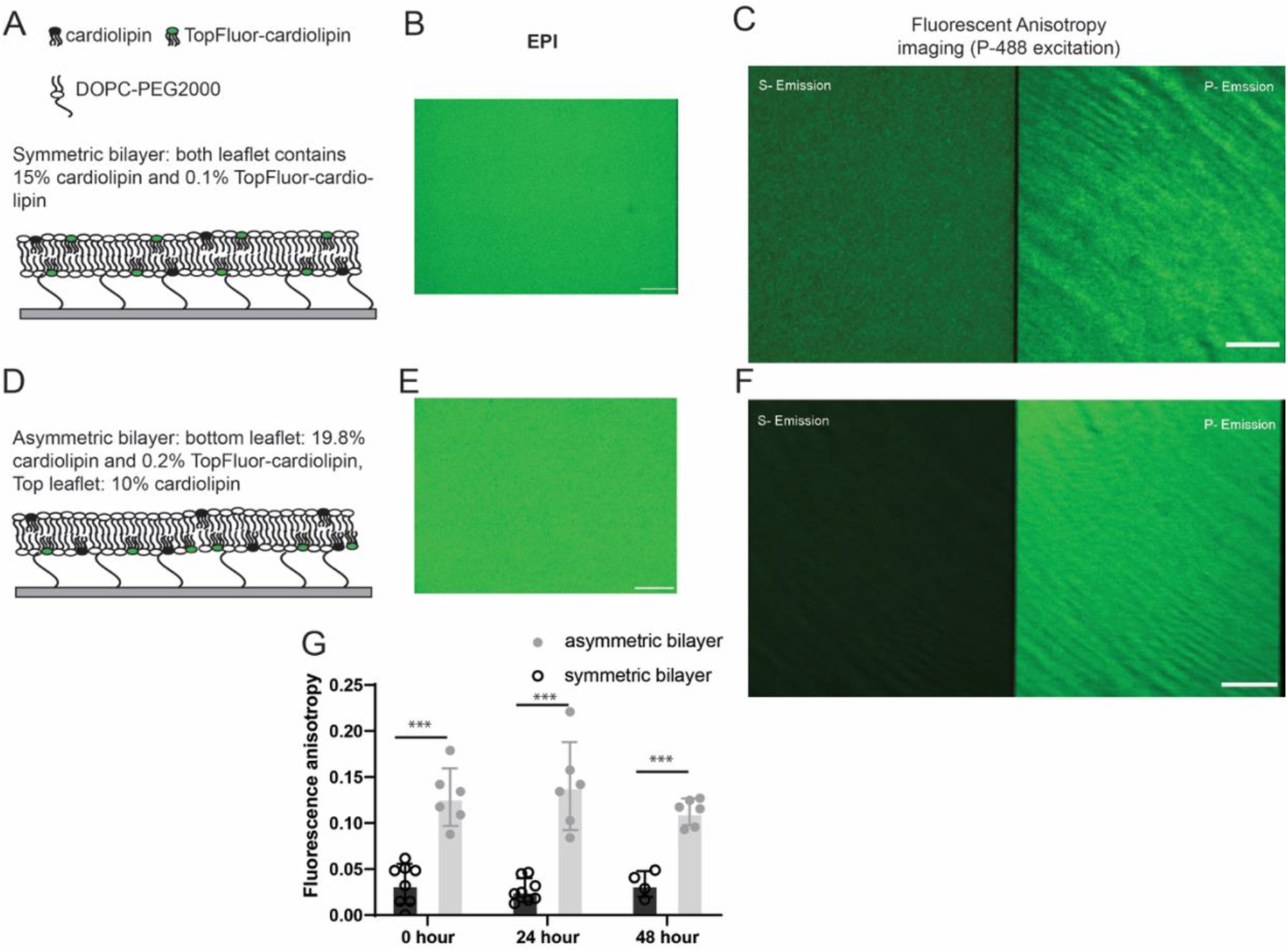
Scheme for symmetric and asymmetric polymer-supported bilayers and evaluation of cardiolipin asymmetry by fluorescence anisotropy. Bilayers with symmetric and asymmetric CL distribution (A and D) show no difference in fluorescence emission intensity under EPI-fluorescence illumination (B and E). However, under polarized excitation (*P*_*polarization*_) the fluorescence intensity of P and S emission channels show significant differences (C and F). Fluorescence anisotropy measurements over time (G) show that bilayer asymmetry was stable over a 48 h time period. Scale bar of each image: 10 μm. Data were collected from 4 bilayers at 5 random locations collected from each bilayer (ns, P > 0.05; *, P ≤ 0.05; **, P ≤ 0.01; ***, P ≤ 0.001; ****, P ≤ 0.0001).

We analyzed TIRF-fluorescence anisotropy images to evaluate lipid asymmetry, as performed previously in cellular membrane and model liposome membranes (Li et al., 2019;St Clair et al., 2020). Both symmetric and asymmetric bilayer were labeled with the same total concentration of fluorophore TopFluor-cardiolipin. EPI-fluorescence images show the same fluorescence intensity (Fig 1B and E). Fluorescence anisotropy of the asymmetric bilayer (average *r*_asy_ = 0.125) was approximately four times higher than that of the symmetric bilayer (average *r*_sym_ = 0.024) (Fig 1C, F and G). Models of plasma membrane asymmetry show high levels of cholesterol and phospholipid scrambling, which causes membranes to homogenize over time (Doktorova et al., 2020). To evaluate the potential for CL flip-flop and loss of bilayer asymmetry anisotropy measurements were also performed after 24 and 48 hours. There was no statistically significant anisotropy change after 48 hours (Fig 1G). Finally, to validate bilayer asymmetry, we also measured anisotropy after reconstitution of l-Opa1, and found similar levels of anisotrophy to those in the absence of protein (Supplemental Figure 1). These findings are consistent with simulation studies showing that an asymmetric distribution of CL is thermodynamically favorable (Kagan et al., 2015; Elias-Wolff et al., 2019).

### l-Opa1 mediated homotypic membrane tethering is dependent on CL distribution and influenced by membrane asymmetry

In our previous studies, we showed CL is important for both l-Opa1 (homotypic and heterotypic) and s-Opa1-mediated membrane tethering (Ge et al., 2020b). l-Opa1 tethering can potentially occur through homotypic protein-protein interactions, or heterotypic protein-lipid interactions. Here, we first investigated homotypic l-Opa1 membrane tethering in a bilayer with asymmetric CL distribution. We compared four conditions varying CL presence in the bilayer and proteoliposome. Initial validation of our system show that we can generate asymmetric bilayers that are stable even with 10% inter-leaflet difference in CL concentration. To evaluate Opa1 function under extreme membrane asymmetry, we completely depleted the CL on the top leaflet in our reconstitution experiments, while leaving 20% CL in the bottom leaflet. Under homotypic l-Opa1 conditions (reconstituted in both bilayer and liposome), the presence of CL does not significantly stimulate GTP-independent membrane tethering, independent of the location of CL or leaflet distribution (Fig 2 A-ii apo and A-iii apo compared to A-iv).

**Fig 2.**
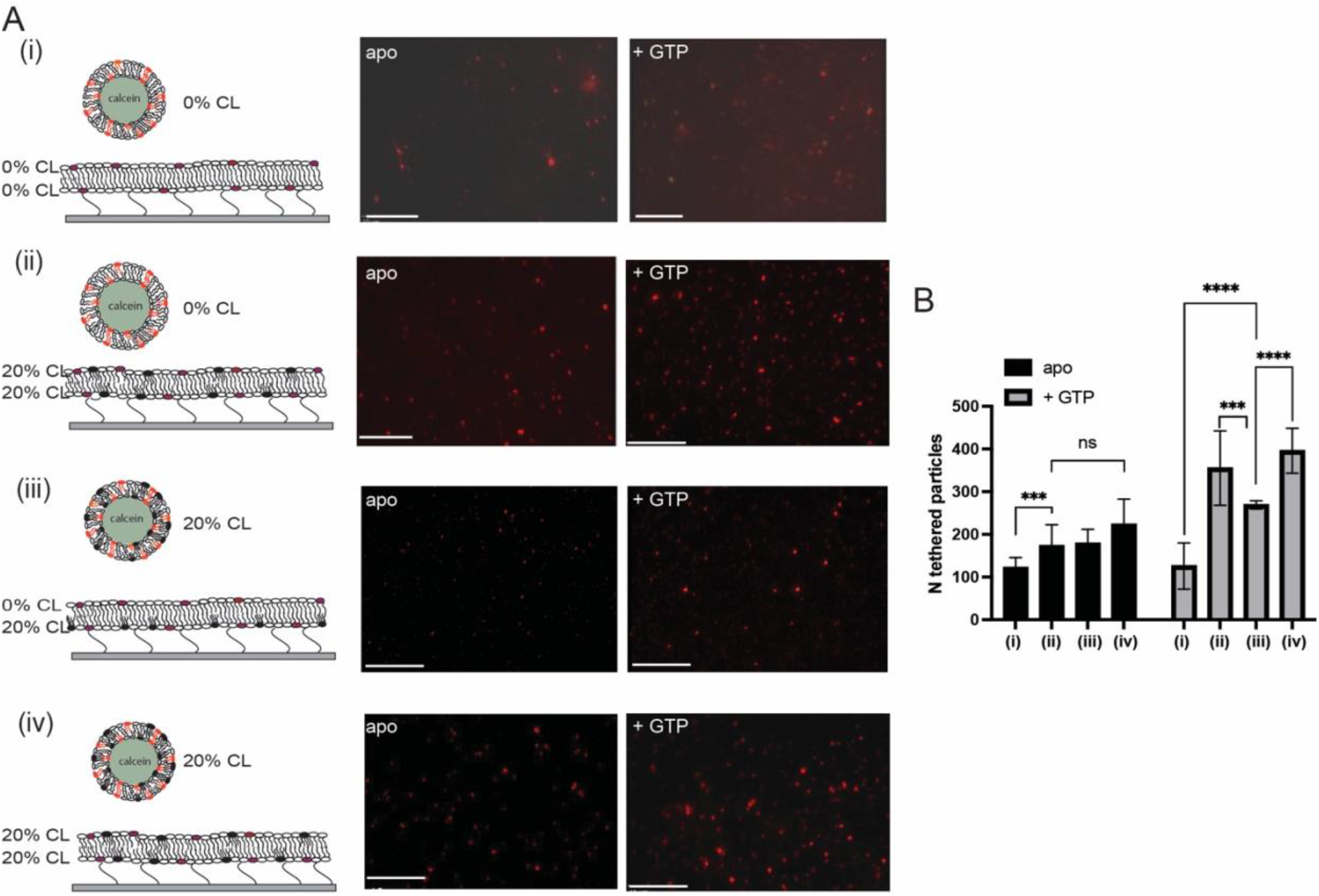
Effect of cardiolipin distribution and membrane asymmetry on l-Opa1 mediated membrane tethering. We compared four conditions: no CL in the bilayer or proteoliposomes (Fig 2A i); 20 % (mol) CL symmetrically distributed in both leaflets of the supported bilayer, no CL in liposomes (Fig 2A ii); no CL in only the top leaflet of the bilayer, proteoliposomes with symmetric 20% CL (Fig 2A iii); 20% CL liposomes and bilayers, symmetrically distributed in both leaflets (Fig 2A iv). Scale bar: 10 μm. Data were collected from 3-5 independent experiments at 5∼10 random positions. Statistical analyses were performed using a t-test. The illustrations indicate lipid distribution in the membrane, with both bilayers containing reconstituted l-Opa1.

Our previous data, and observations from others (Ban et al., 2010), show that membrane embedded CL can mediate nucleotide-dependent membrane tethering. Interestingly, we observe reduced particle tethering when CL was absent from the top leaflet of the planar bilayer, compared to the symmetrically distributed case (Fig 2A iii and iv). In our previous studies, we found that GTP-independent tethering did not depend on the presence of cardiolipin (Ge et al., 2020b). Our quantification here shows that when CL is present in both leaflets of the bilayer (regardless of the CL content in proteoliposomes), addition of GTP induces ∼three-fold increase in proteoliposome tethering to the bilayer (Fig 2B, 2A ii and iv). In contrast, when CL is depleted from only the outer leaflet of the bilayer, GTP triggers ∼1.5 fold increase in proteoliposome tethering to the bilayer (Fig 2 A iii, 2B).

### s-Opa1 promotes membrane tethering in asymmetric CL bilayers

Proteolytic processing of the transmembrane anchored l-Opa1 to s-Opa1 is a key cellular regulator of mitochondrial inner membrane fusion and cristae state (Del Dotto et al., 2017;Wang et al., 2021) Our previous *in vitro* reconstitution studies using symmetric membranes, showed that stoichiometric levels of s-Opa1 were optimal for stimulating fast and efficient membrane fusion by l-Opa1 (Ge et al., 2020b). This result reconciled different cellular observations of mitochondrial reticulum phenotypes when modulating the fusion-fission equilibrium (Song et al., 2007; Ehses et al., 2009; Anand et al., 2014; Wang et al., 2021). Here, we investigate the effect of s-Opa1 addition in an asymmetric CL bilayer system. Unexpectedly, depleting CL from the top leaflet of the supported planar bilayer changed the effect of s-Opa1 on proteoliposome tethering.

Upon addition of s-Opa1 at a sub-stoichiometric ratio of 1:10 s-Opa1:l-Opa1, the number of tethered proteoliposomes increased >5 fold, in the presence of GTP (Fig 3B). Addition of stoichiometric levels of s-Opa1 to l-Opa1, did not increase proteoliposome tethering to the bilayer. Instead, lipid tubes were observed to form. These tubes were tethered to the bilayer. When s-Opa1 levels were in excess of l-Opa1, fewer proteoliposomes tethered to the bilayer, and the lipid tubes dissociated from the bilayer surface (Fig 3D and E).

**Fig 3.**
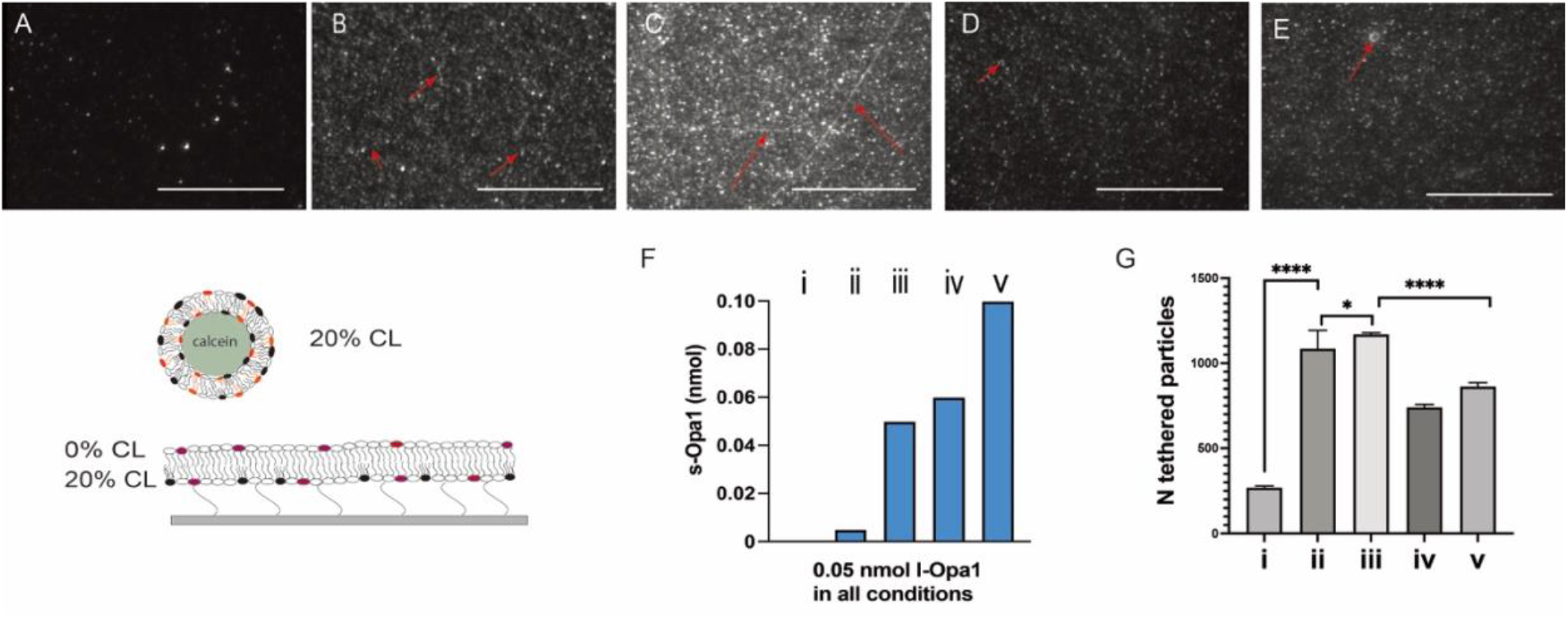
Effect of s-Opa1 on proteoliposome tethering to an asymmetric CL bilayer. A. In the presence of GTP, few l-Opa1 proteoliposomes are tethered to an asymmetric CL bilayer containing l-Opa1. Upon s-Opa1 addition, proteoliposome tethering increases significantly. Panels B-E show representative images varying l-Opa1 and s-Opa1 concentration shown in panel F as i-iv. l-Opa1 concentration was fixed at 0.05 nmol in all conditions. Scale bar, 20 μm. Red arrows indicate tubular structures tethered to the bilayer. G. Quantification of the number of proteoliposomes tethered to the bilayer, based on 10-20 random points in each experiment, for a total of 3-5 different experiments. Data were analyzed using t-test. The illustrations indicate lipid distribution in the membrane, with both bilayers containing reconstituted l-Opa1.

### Cardiolipin in the outer leaflet plays roles in efficient l-Opa1-mediated homotypic fusion and s-Opa1 stimulated fusion

Model membrane and cellular studies show l-Opa1 is required for membrane fusion, and s-Opa1 plays an important stimulatory role (Song et al., 2007; Ge et al., 2020b). Previously, we developed a three color imaging assay to evaluate the kinetics of membrane fusion (Ge et al., 2020b). The fluorophores used for monitoring fusion differ from those we used for characterizing anisotropy. Briefly, membrane docking was defined by the FRET signal between Cy-5 (bilayer label) and TexsasRed (proteoliposome label). Lipid demixing or hemifusion events were reported by a proteoliposome marker (Texas Red) diffusing into the lipid bilayer. Release of calcein from the proteoliposomes (loaded at quenched concentrations) reported on pore opening events and their kinetics. Using this assay, we observed low levels of hemifusion and full fusion (pore opening) in a homotypic l-Opa1 format (l-Opa1 reconstituted into the supported bilayer and proteoliposomes, both with symmetric CL distribution). Under these conditions, addition of s-Opa1 dramatically stimulated full fusion (Ge et al., 2020b).

Here, in l-Opa1 reconstituted bilayers with asymmetric CL distribution, we observed fewer hemifusion and full fusion events (Figure 4D and G, respectively), compared to our previous finding in symmetric bilayers (Ge et al., 2020b), where 20% of the total particles hemifuse with the bilayer. Under asymmetric conditions, addition of s-Opa1 did not increase the efficiency of membrane hemifusion or fusion (Figure 4D, G).

**Fig. 4.**
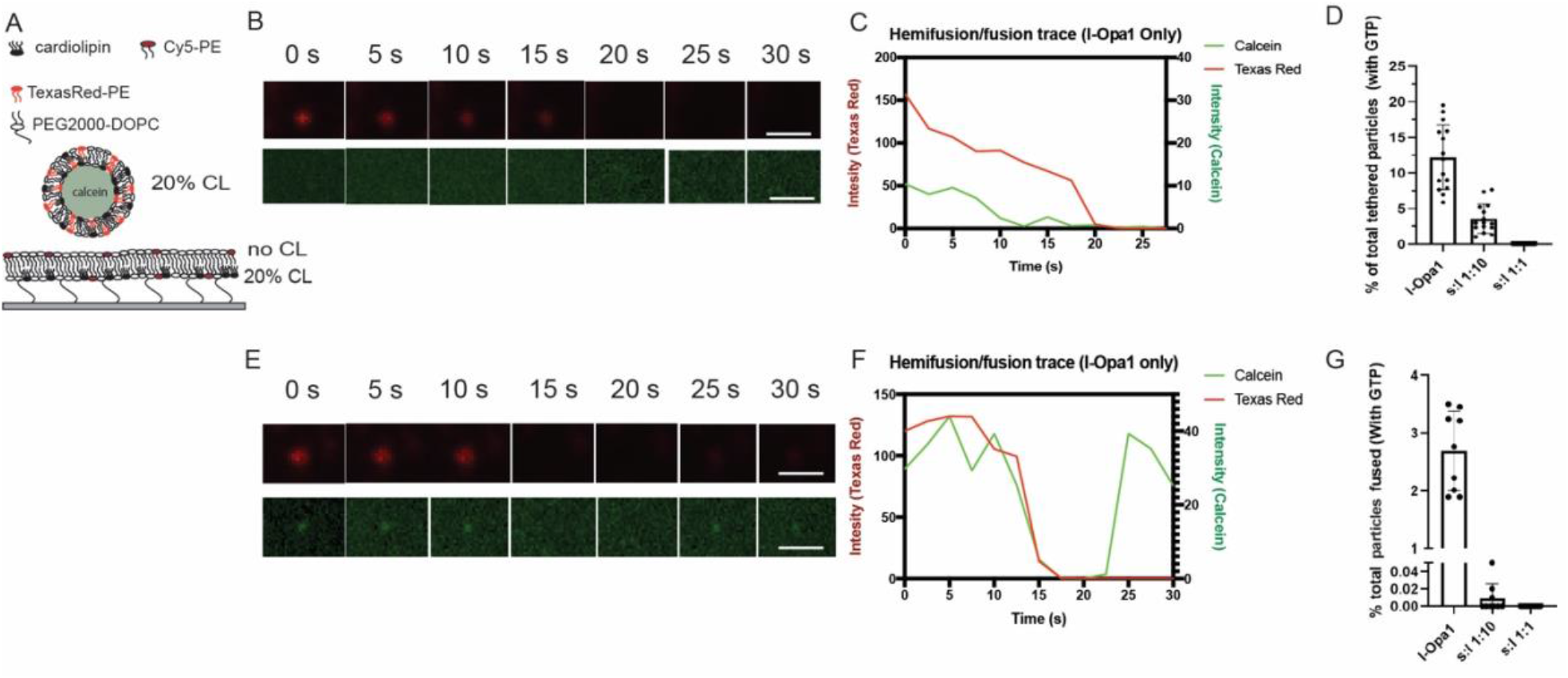
CL distribution determines the efficiency and kinetics of l-Opa1 mediated membrane lipid demixing (hemifusion) and content release. A. Schematic of the experiment showing asymmetric cardiolipin distribution in the membrane, both bilayers with l-Opa1 reconstituted. The lipid bilayer top leaflet was CL free, while the bottom leaflet contained 20 %(mol) CL. B. Example of hemifusion event without calcein release. The total process takes ∼20 s. Full fusion kinetics shown in E, with calcein release observed after 30 seconds of total recording time. The total number of particles analyzed from 3∼5 bilayers quantified in D (hemifusion) and G (full pore opening), under the asymmetric cardiolipin distribution conditions illustrated in A. Data were collected from 3 different experiments. Scale bar in images, 1 μm.

We also observed a significant delay in lipid demixing with l-Opa1 alone. Compared to symmetric bilayers, the average time for lipid demixing in asymmetric bilayers changed from 15 s to 20 s (in the ∼50 events calculated in 3 different experiments) (Figure 4B, C), whereas the content release time changed from 20 s to 30 s or more (Figure 4E, F). The calcein release time was also significantly longer (from milliseconds to several seconds) with asymmetric CL bilayers, possibly suggesting an effect on calcein diffusion due to slow fusion pore expansion. Together these observations all point to the importance of CL in the outer, IMS-facing leaflet in efficient s-Opa1 stimulated fusion.

## Discussion

The mitochondrial inner membrane (IMM) is expected to have an asymmetric distribution of CL under physiological conditions (Kagan et al., 2014; Pizzuto and Pelegrin, 2020). In this study, we used an *in vitro* model IMM to explore Opa1 activity upon selectively depleting CL from the outer leaflet. We found three interesting effects under these asymmetric CL conditions. First, removing CL from the outer leaflet reduces GTP-dependent homotypic l-Opa1 tethering. Second, we observe lipid tube formation when sub-stoichiometric levels of s-Opa1 are added l-Opa1 tethered proteoliposomes containing CL. Third, asymmetric CL distribution slows pore opening kinetics. These observations all point to the importance of CL in the outer leaflet in mediating efficient fusion.

Recent structures of s-Opa1 and its yeast orthologue s-Mgm provide candidate interfaces for understanding self-assembly. These structures all show Opa1 has a canonical dynamin-family GTPase domain arrangement, with GTPase domain, bundle signaling element (BSE), stalk domain and paddle region (Faelber et al., 2019; Zhang et al., 2020). Basic and hydrophobic residues in the paddle region directly interact with membranes containing CL (Faelber et al., 2019; Yan et al., 2020; Zhang et al., 2020). Mutation of these residues results in decreased affinity of s-Opa1 to the lipid bilayer and affects GTPase activity (Zhang et al., 2020). In Cryo-ET studies, interfaces formed by the helical stalk region mediate self-assembly (Faelber et al., 2019).

Our observation of reduced tethering under asymmetric conditions suggests that the presence of CL in the outer leaflet facilities inter-membrane interactions. We still lack high-resolution structural information for l-Opa1. Existing observations suggest potential modes of CL-influenced tethering may be at play. CL in the bilayer might stabilize a conformation of Opa1 such that it can interact in a homotypic fashion with another copy of Opa1 in the proteoliposome bilayer. It should be noted, that although the outer leaflet of the bilayer was fabricated without CL for our assay, recombinant l-Opa1 was purified in the presence of CL. We estimate that during reconstitution, around 10^−12^ mol of CL may be reconstituted into the outer leaflet, resulting in an outer leaflet with 0.01 %(mol) CL, which is higher than the total amount of l-Opa1 reconstituted in the planar bilayer. Based on previous GTP-assays, however, this level of CL is unlikely to significantly affect Opa1 activity (Ge et al., 2020b).

Membrane tubes are an intriguing *in vitro* observation of Opa1 behavior, with several candidate *in vivo* roles proposed. In our previous work with symmetric bilayers, we did not observe lipid tube formation with the protein concentrations used under three different conditions, namely, l-Opa1 or s-Opa1 alone, and l-Opa1 mixed with s-Opa1 (Ge et al., 2020b). Interestingly, we do observe tube formation with asymmetric bilayers lacking CL in the outer leaflet, upon addition of s-Opa1. With a bilayer containing symmetric inter-leaflet CL distribution, no lipid tubules were observed previously. In this work (and our previous studies), we used low concentrations of reconstituted protein combined in the bilayers and liposomes, making the total l-Opa1 concentration of 6.5×10^−12^ mol/ml (protein:lipid, 1:5000), and a final s-Opa1 concentration of 6.5×10^−11^ mol/ml, (final protein:lipid 1:500) (Ge et al., 2020b). The Daumke and Sun groups determined helical assemblies of s-Mgm1 and s-Opa1, respectively, decorating the outside of lipid tubes and inducing positive curvature (Faelber et al., 2019;Zhang et al., 2020). Faelber and colleagues also observe s-Mgm1 tubes decorating the interior of tubes, with negative curvature (analogous to the topology expected within cristae). Li and co-workers suggest a s-Mgm1 trimer may help ‘bud’ regions of bilayer, through puckering membranes with positive curvature (Yan et al., 2020). In these reports, s-Opa1 was added at 1:1 (protein: lipid molar) ratio, for a total concentration of s-Opa1 5×10^−9^ mol/ml (Ban et al., 2010; Zhang et al., 2020). In these studies, interaction of s-Opa1 with CL is important for tubulation. In our asymmetric CL experiments, we observe tubulation at even lower concentrations. Due to the lack of CL on the outer (IMS facing) leaflet, we suspect that the major source of membrane for generating the tubes is the liposomes. Whether aspects of tubulation represent an intermediate in fusion will require further investigation. Future studies with asymmetric liposomes will also help discriminate s-Opa1 binding modes. In the asymmetric CL setting, s-Opa1 may interact with l-Opa1, and this heterotypic interaction might explain the observed increase in tethering.

In our previous studies under symmetric CL conditions, we observed fast and efficient membrane fusion upon addition of stoichiometric levels of s-Opa1 (Ge et al., 2020b). In this study, asymmetric CL distribution, specifically the lack of CL in the outer leaflet, appears to stall progression to fusion, resulting in the growth of tubes. Furthermore, addition of s-Opa1 dissociated tubes from the bilayer, suggesting a hetero-oligomeric l-Opa1:s-Opa1 interaction plays a role in association of these tubes with the supported bilayer. We did not observe any hemifusion events between tubes and the supported bilayer, emphasizing the importance of CL in this leaflet for proceeding through fusion. The lack of stimulatory effect upon addition of s-Opa1 to a homotypic l-Opa1 arrangement also suggests s-Opa1 assembly (or interaction) with the supported bilayer plays a role in fusion.

The *in vitro* system we describe here opens exploration of how membrane lipid composition and distribution influence the interplay of different physical properties to regulate fusion and ultrastructure. For example, fusion efficiency may be influenced by the combination of membrane elastic properties, including monolayer spontaneous curvature and bending modulus. (Fan et al., 2016). Calculations based on physical models suggest spontaneous curvature is a factor regulating the elastic energy of the pre-fusion planar bilayer. In experiments performed here, we investigated supported lipid bilayers with cardiolipin depleted from the outer leaflet. If the unsaturated, four acyl-chain CL is depleted from the outer leaflet, intrinsic negative curvature may be disfavored. Negative spontaneous curvature would be expected to favor the formation of hemifusion stalk, and be important at regions such as the cristae tip. The converse situation, with cardiolipin absent from the inner-leaflet will not only eliminate Opa1-lipid interactions, but also potentially disfavor positive curvature. Lowering the barrier to positive curvature may important at regions like the cristae junction. Depletion of CL in either leaflet could result in bilayers with a higher bending modulus than that of a symmetric bilayers. Changes in the bending modulus could increase the energy barrier from planar membrane to a highly curved states. Other physical properties likely influenced by cardiolipin distribution include line tension, hydrophobic mismatch and membrane packing. Investigating how these factors regulate membrane rearrangement are of great interest for future studies.

## Conclusion

In this study we tested the effects of selectively removing CL from one leaflet of a model of the mitochondrial innermembrane. We observed that simply removing CL from the IMS facing leaflet resulted in interesting changes in membrane tethering, and stalled fusion. Furthermore, we observe formation of membrane tubes under these conditions. Low levels of CL are expected to be present in the IMS-facing leaflet under physiological conditions. These observations point to important conformations and assembly states of Opa1 regulated by the presence of CL. These studies also suggest that interesting membrane properties resulting from lipid asymmetry remain to be explored. The specific protein arrangements and membrane states of fusion are key questions for future integrated structural and biophysical studies.

## Acknowledgements

We thank all members of the Chao Lab for helpful discussions and suggestions. This work was supported by funding from the Charles H Hood Foundation Child Health Research Award and the National Institutes of Health (R35GM142553) to L.H.C.

---

**Supplemental Fig. 1.**
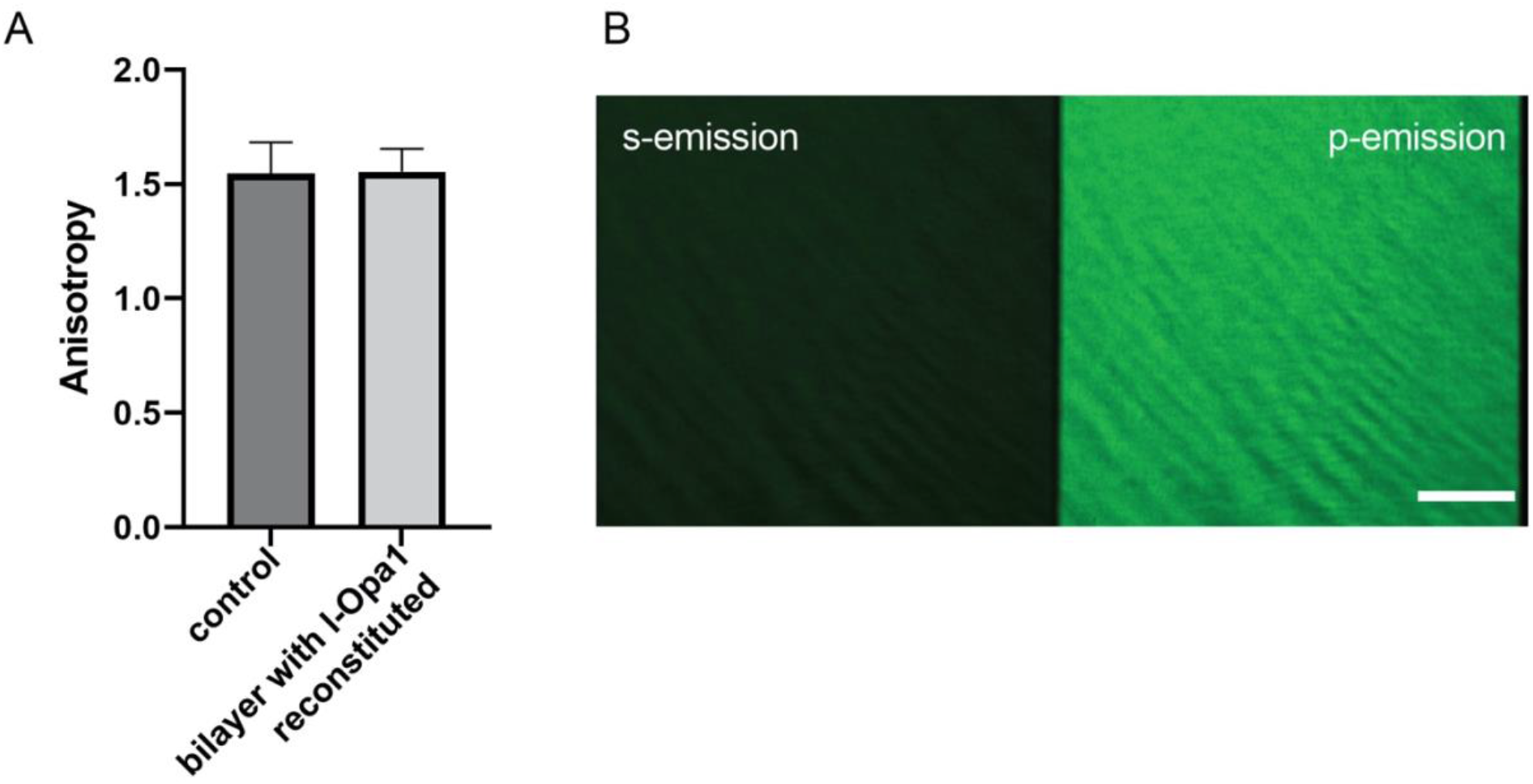
l-Opa1 reconstitution does not affect CL asymmetry. A. Calculated anisotropy before and after l-Opa1 reconstitution suggest a smiliar cadiolipin distribution in the bilayer. B. shows the fluorescence images obtained in s-emission and p-emission channels.

